# Nanoscale interaction of endonuclease APE-1 with DNA characterized by atomic force microscopy

**DOI:** 10.1101/2024.02.11.579811

**Authors:** Sridhar Vemulapalli, Mohtadin Hashemi, Yinglink Chen, Suravi Pramanik, Kishor K. Bhakat, Yuri L. Lyubchenko

**Author notes:** Corresponding authors Yuri L. Lyubchenko, Kishor K. Bhakat.

## Abstract

Apurinic/apyrimidinic endonuclease 1 (APE1) is involved in DNA replication, repair, and transcriptional regulation mechanisms. This multifunctional activity of APE1 should be supported by specific structural properties of APE1 that have not yet been elucidated. Here we applied atomic force microscopy (AFM) to characterize the interactions of APE1 with DNA. Complexes of APE1 with DNA containing G-rich segments were visualized, and analysis of the complexes revealed the affinity of APE1 to G-rich DNA sequences. Furthermore, loops in the DNA-APE1 complexes were visualized, and their yield was as high as 53 %. However, the loops were non-specific, with quantitative analysis revealing the yield of loops bridging two G-rich DNA segments to be 41%. Analysis of protein size in various complexes was performed, and these data showed that loops are formed by APE1 monomer, suggesting that APE1 has two DNA binding sites. The data lead us to a model for the interaction of APE1 with DNA that describes its molecular site search mechanism. The new properties of APE1 in organizing DNA, by bringing two distant sites together, may be important for facilitating the scanning for damage and coordinating repair and transcription.

## 1 Introduction

Apurinic/apyrimidinic endonuclease 1 (APE1) is a multifunctional protein involved in DNA repair, specifically base excision repair (BER) and nucleotide incision repair (NIR) (1-4). APE1 is essential for maintaining genomic stability by repairing the DNA damage caused by reactive oxidation species, alkylating agents, and ionizing radiation (4-6). APE1 cleaves abasic sites in the DNA, which occur due to damage or as a repair intermediate (1,2,5,7,8). In addition to its role in DNA repair, APE1 interacts with transcription factors such as p53, NF-κB, and HIF-1α and plays a vital role in DNA replication and is involved in resolving stalled replication forks by repairing the damaged DNA during DNA replication, repair, and recombination processes (3,5,6,9-11). It has been reported that APE1 interacts with several key enzymes like DNA polymerase delta and stimulates the activity of flap endonuclease 1 (FEN1) (6,10,12,13).

APE1 directly interacts with and binds G4 structures formed by G-rich DNA sequences, suggesting its role in regulating gene expression (1,2,14-16). In addition, APE1 has been reported to regulate the stability and dynamics of G4 structures by nicking or cleaving the DNA backbone at specific positions, which affects the folding and unfolding of the G4 structure (1,2,5,11). APE1 interacts with other G4-binding proteins, such as nucleolin and hnRNPU, and regulates their binding to G4 structures (1,2,16,17). APE1 interactions with G4 structures and binding proteins offer potential therapeutic targets for cancer treatment and other diseases (1-3,5,7-11,18-21).

Participation of APE1 in transcription regulation led to the hypothesis of APE1 being able to bring together two specific DNA segments, which, in case the two segments are on the same DNA molecules, can lead to looping. Enrichment of APE1 in gene regulatory regions binds to G4 structures, and participation in transcriptional regulation led to the hypothesis that APE1 can bring together two specific DNA segments on the same DNA molecules, forming a loop. However, there is no direct evidence of APE1-mediated DNA looping. The formation of DNA loops requires binding APE1 to two DNA segments. Loop formation can be achieved by the interaction of two monomers bound to two sites or by binding two DNA segments by one monomer, suggesting that two DNA binding sites are present in the APE1 monomer.

We address these questions using AFM to characterize the APE1-DNA complexes directly. We have previously shown that AFM is instrumental in imaging various protein-DNA complexes, reviewed in (22). Specifically, we characterized looped protein-DNA complexes, such as those formed by restriction enzymes (23,24). Importantly, using AFM, we identified additional DNA binding sites in EcoRII endonuclease, allowing the formation of double-looped complexes (25). Here we applied AFM to characterize the interaction of APE1 with DNA using a DNA substrate containing two well-separated G-rich segments. Using this approach, we demonstrate the affinity of APE1 to G-specific motifs. The formation of loops was also demonstrated, but in addition to specific loops between the G-rich segments, non-specific loops are also formed. However, no G-quadruplex structures were identified on the DNA substrate alone, suggesting that their formation is not required for APE1-specific binding and such structures can be stabilized by APE1 binding. Finally, loops are formed by the monomeric APE1 protein, suggesting that the protein has two DNA binding sites.

## 2 Experimental procedure

### 2.1 Materials

#### 2.1.1 APE1-protein

The full-length APE1 coding sequence was inserted in the pET15b vector (Novagen) at NdeI/Xho I sites for expression of APE1 in *E. coli* Rosetta 2 strain. The DNA sequence of the APE1 was confirmed by UNMC genomic core. APE1 protein was purified as previously described (26) with slight modifications. After transforming with the pET15b-based APE1 expression plasmid, *E. coli* were grown to 0.6 OD at 600nm. APE1 expression was then induced with 0.5 mM isopropyl-β-D-thiogalactopyranoside (IPTG) at 18° C for 16 h. The cells were then suspended in a buffer containing 20 mM Tris (pH 8.0) and 0.5 M NaCl, sonicated, and centrifuged. The supernatant was loaded onto Ni-NTA (Qiagen) column (3 mL), run, and then eluted with buffer containing 200 mM imidazole. The eluate was dialyzed against 20 mM Tris-Cl (pH 8.0), 100 mM NaCl, 1 mM EDTA, 1 mM dithiothreitol (DTT), and 10% glycerol. The poly His-tag in the protein was cleaved by overnight incubation at 4° C with thrombin. The APE1 was finally purified by FPLC using an SP-Sepharose column (LCC-500 PLUS; Pharmacia), and the final preparation was dialyzed against 20 mM Tris (pH 8.0), 300 mM NaCl, 0.1 mM EDTA, 1 mM DTT, 50% glycerol, and stored at −20° C.

#### 2.1.2 DNA Substrates

A 673 bp DNA segment of *c-MYC* gene promoter (−25 to -648 bp with respect to the transcription start site) was amplified by PCR, and formed the DNA substrate containing two G-rich motifs (Fig. 1A). For the PCR reaction, 100ng of human genome DNA and PfuUltra High-Fidelity DNA polymerase (#600380) were used with the primers: Forward primer: AGGGTTTGAGAGGGAGCAAAAG; Reverse primer: CTCGGGTGTTGTAAGTTCCAG. Similarly, DNA without G-rich motifs with a length of 612 bp, as shown in (Fig.1A), was obtained by performing PCR of the plasmid.

**Figure 1.**
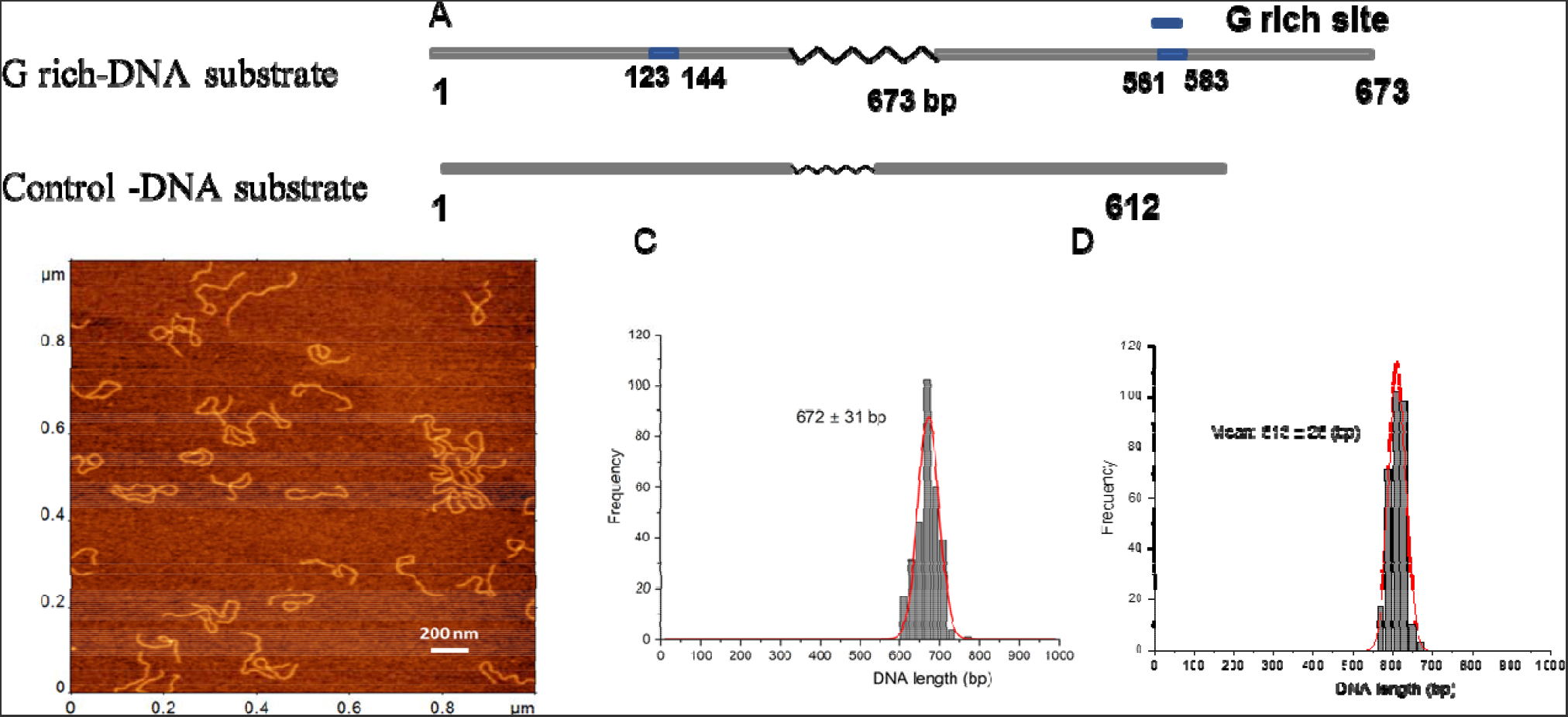
DNA substrates, AFM image, and contour length. (A) The schematic for the G rich-substrates (upper scheme) and the control (bottom). G-rich motifs with and 22 bp long are located at 123-144 bp G-rich site at 561-583 bp and shown in blue. The non-G-rich DNA substrate with 612 bp in length was used as a control. (B). A typical 1x1 μm AFM scan of G-rich DNA substrate (C) and (D) are histograms for the contour length measurements for G-rich DNA substrate and the control respectively. Each distribution is approximated with single Gaussians built with a bin size of 20 bp. The contour length values in base pairs and standard deviations are indicated for each histogram.

Both DNA substrates were gel-purified as described (23). Briefly, the PCR product was run on a 1% agarose gel. The product bands corresponding to the expected length of the DNA were excised, and DNA was extracted and purified using the Qiagen DNA gel extraction kit (Qiagen Inc., Valencia, CA). The final DNA concentration was determined by absorbance at 260 nm using a NanoDrop spectrophotometer (NanoDrop Technologies, Wilmington, DE).

#### 2.1.3 APE1-DNA synaptosome assembly

DNA was mixed with APE1 enzyme at the molar ratio 1:1 in 50 mM Tris-HCl buffer containing 50 mM KCl, 2 mM MgCl_2_ with a total volume of 10 μl, and with the final concentrations of DNA and APE1 at 1nM. A reaction mixture for APE1-DNA assembly consisted of a final volume of 10ul with 7ul of 1X buffer A [50 mM Tris HCl (pH 7.5), 50 mM KCl, 2mM MgCl_2_, 0.1 mM EDTA], 1μl of 10 mM DTT, 1μl of DNA, and 1μL of protein. The reaction mixture was incubated for 15 min at room temperature.

#### 2.1.4 Atomic Force Microscopy

The APE1-DNA complexes were deposited on functionalized mica, functionalized with 1-(3-aminopropyl)-silatrane, as described previously (23,24,27). Briefly, 3-4 μl of APE1-DNA reaction mixture was deposited on the functionalized mica surface, incubated for 2 minutes, rinsed with deionized water, dried with argon, and stored under vacuum until imaged.

A typical AFM image scanned 3x3 μm area with 1536 pixels/line under ambient conditions. Imaging was performed with a MultiMode 8 AFM system using TESPA probes (Bruker Nano, Camarillo, CA, USA).

#### 2.1.5 Data Analysis

The contour length of the bare DNA, the APE1-DNA complexes, and the looped the APE1-DNA complexes were measured using FemtoScan software (Advanced Technologies Center, Moscow, Russia) as described previously (23,24), which allows reliable tracing of DNA, as shown in Figure S1. Figures S2 and S3 illustrate the measurements of the protein position and loop size, respectively.

#### 2.1.6 Height and volume of APE1

Grain analysis was performed to measure the height and volume of the free APE1, APE1 complexed with DNA, and APE1 in looped complexes.

## 3 Results

### 3.1. DNA design: Preparation of the substrate with two APE1 sites

We used a 673 bp DNA substrate from the human genome containing two 22 bp long G-rich motifs of the *c-MYC* gene regulatory region. G-rich segments were located at positions 123 bp and 583 bp of the DNA substrate (Fig. 1A). The DNA was selected based on the previous biochemical studies conducted to characterize the APE1 interactions (1,2). The two G-rich sites on the DNA are separated by 417 bp, which according to our previous publications is appropriate for the assembly and the AFM visualization of the protein-mediated DNA loops (23,24).

Typical AFM images of the G-rich DNA substrate are shown in Figure 1B in which DNA appears as smooth filaments. The contour length measurements are shown in Figure 1C. A total of N = 300 particles was analyzed, and a single peak Gaussian function approximation of the histogram gives a mean of 672 ± 31 bp (SD). Similar contour length measurements of the control DNA of 612 bp with no G-rich segments are shown in Figure 1D.

### 3.2. APE1 complex assembly and loop size analysis

The APE1-DNA complexes were assembled at a 1:1 protein:DNA ratio and prepared for AFM imaging. AFM images of the APE1-DNA complexes are shown in Figure 2A with few zoomed images shown in Figure 2B, C. Three different morphologies were identified: bare DNA (Fig. 2B1); DNA with APE1 as bright globular features (Fig. 2B2, 3)) and looped DNA with APE1 as globular features (Fig. 2C1, 2 & 3). The overall yield of complexes is 53%, with the partition of looped and non-looped complexes 22% and 31%, respectively (Table 1).

**Table 1.** Yield of APE1-DNA complexes formed on G rich and non G rich substrate.

| Substrate | N=500 | Ape1-DNA complex<br>(non-looped<br>complexes) | Ape1-DNA complex<br>(complexes with<br>loops) |
| --- | --- | --- | --- |
| G rich-DNA | Yield [%]. | 31 % | 22 % |
| Non-G rich DNA | Yield [%] | 15 % | 4 % |

**Figure 2.**
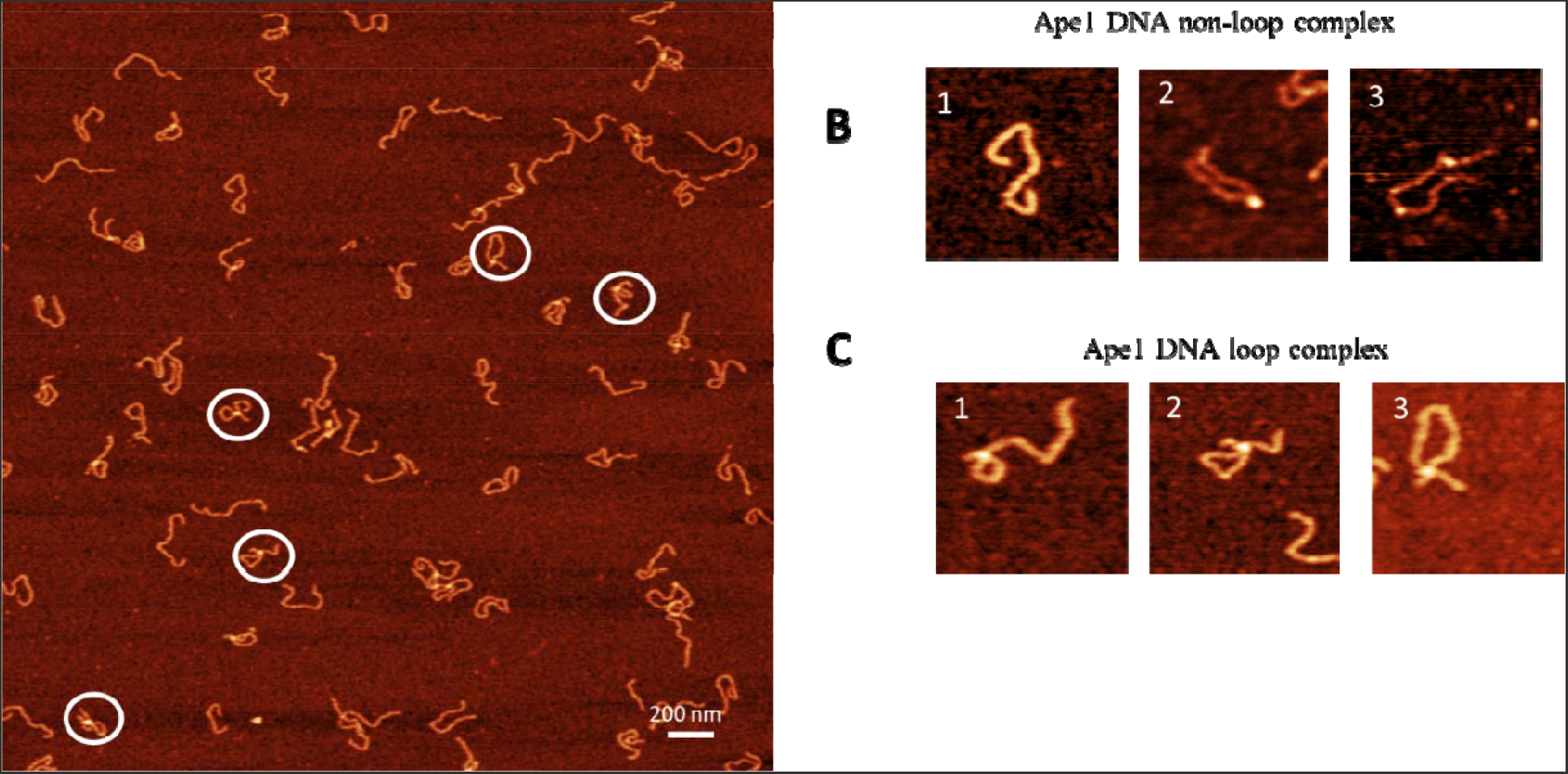
AFM image of complexes of APE1 with G rich DNA complexes (1:1). (A) The AFM image with looped complexes of APE1-G rich-DNA. Zoomed images of complexes circled in (A) are indicated in B and C. (B) A set of images with no APE1 bound – frame 1 and non-pooed complexes with one APE1 bound (frame 2) and two APE1 bound (frame 3). (C) A set of three looped complexes with different sizes of loops.

Similar data were obtained for the control DNA substrate containing no G-rich sequences. The AFM images are shown in Figure 3A. Similar to the G-rich DNA substrate, three different morphologies were observed with selected zoomed images shown in Figure 3B, C. These are free DNA (Fig. 3B1), DNA with bright features (Fig. 3B2, 3), and looped complexes (Fig. 3C1, 2 & 3). The overall yield of complexes is 19 %, with yield of looped complexes being 4% (Table 1).

**Figure 3.**
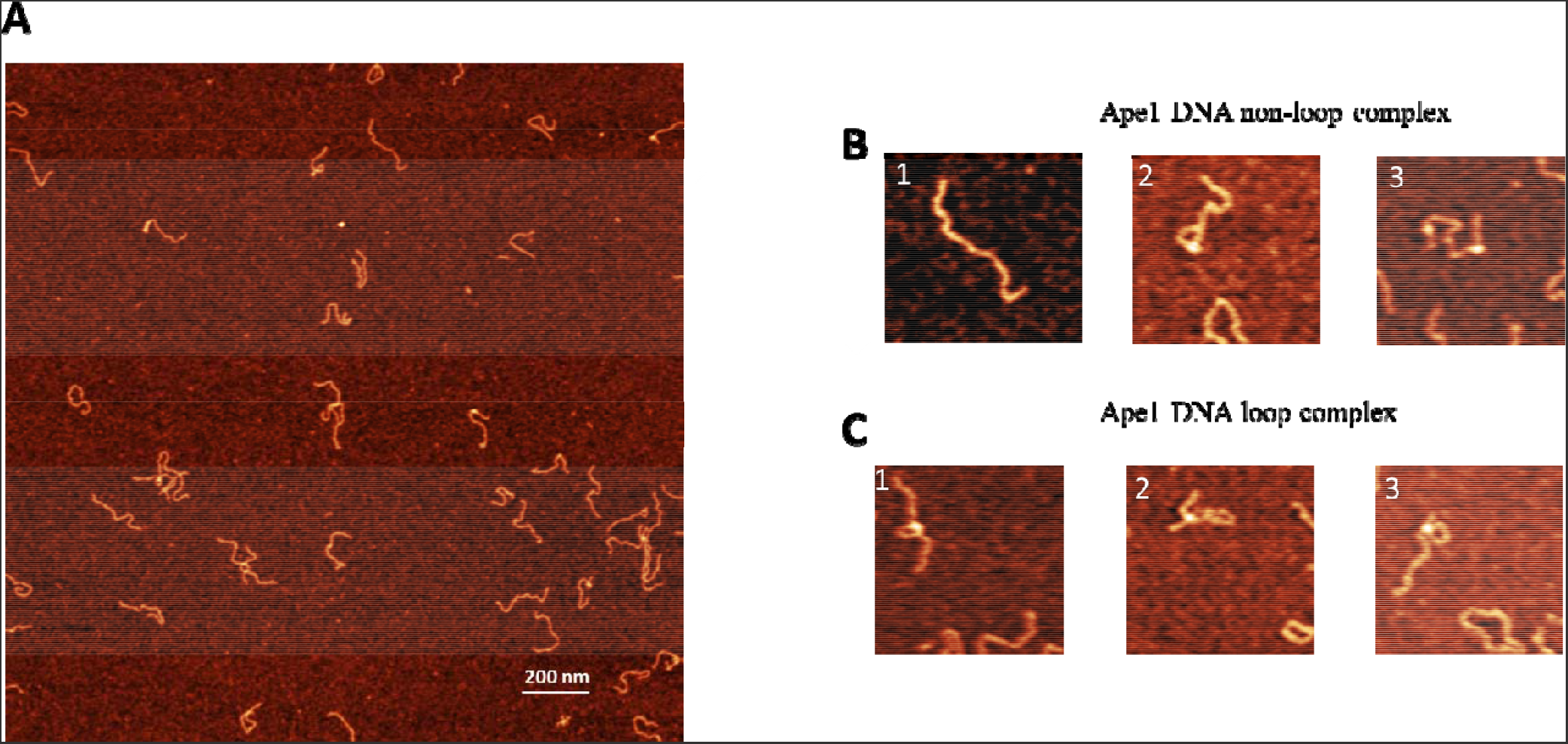
AFM image of complexes of APE1 with a non-G rich DNA complexes (control substrate). (A) A typical AFM scan with 3x3 in size. shows the AFM image with looped complexes of APE1-Non-G rich-DNA. (B) and (C) show a few examples of complexes with liear morphology and looped DNA complexes, respectively.

### 3.3. AFM data analysis: Positioning of APE1 on DNA

Given the relative symmetry in the position of G-rich segments on the DNA, relative to the DNA ends (123-144 bp and 561-583bp), we mapped positions of APE1 on the G-rich DNA by measuring the length of the distance between the bright features and closest DNA end (Fig. 4A). The measurements were made for 300 complexes and the results are shown as a histogram in Figure 4B. Green vertical lines indicate positions of the G-rich motifs in the DNA molecule. Positions of APE1 within the range of the green lines are considered as specific interactions of APE1 with DNA.

**Figure 4.**
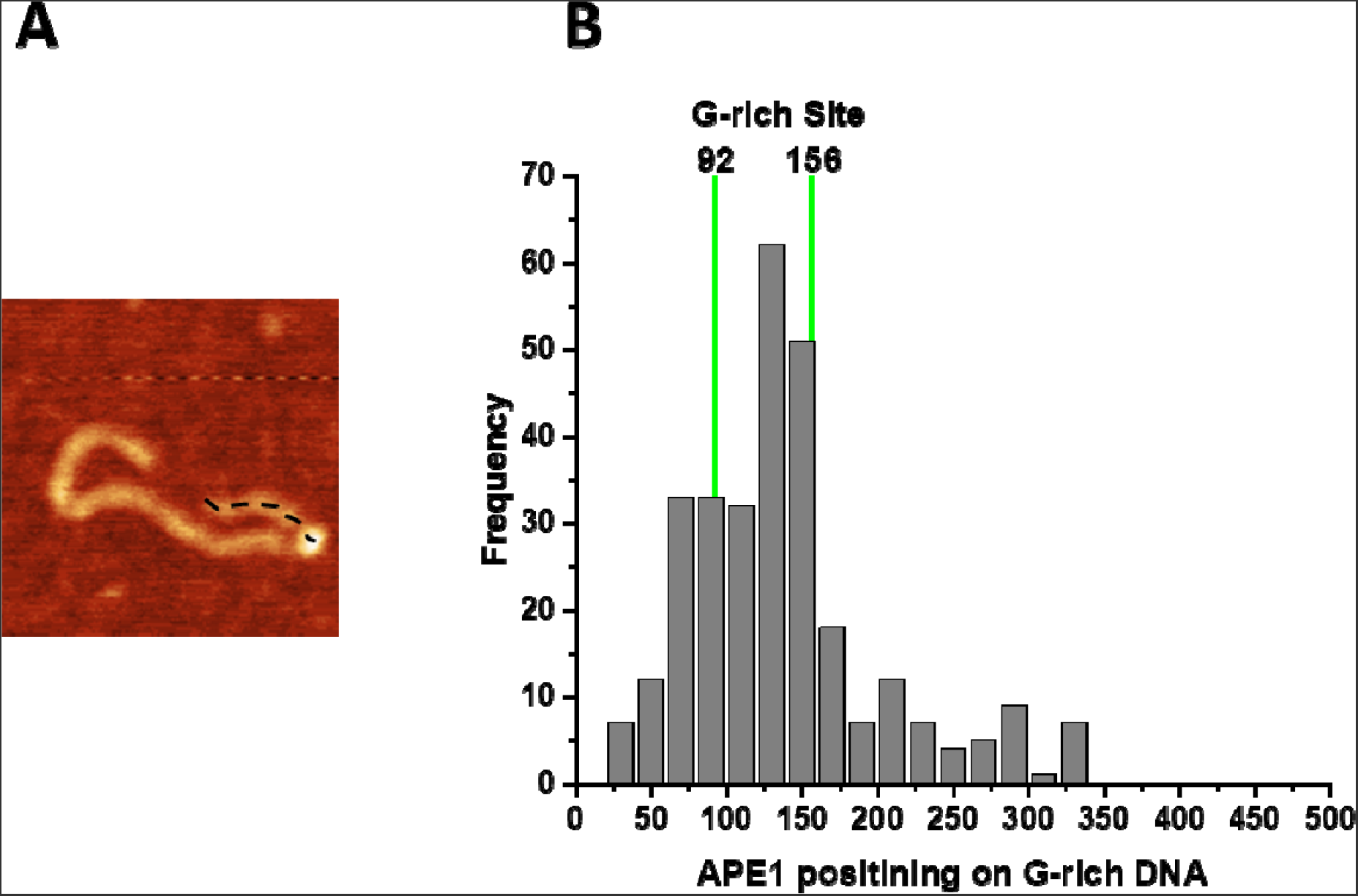
Mapping of the APE1 positions on the G rich-DNA substrate. (A) AFM image of APE1-G rich-DNA complex. Dotted line illustrates the contour length of the short arm measured from the DNA end to the center of the protein. (B) The histogram of APE1 mapping performed over 300 molecules. Vertical green lines correspond to the range so distances from both DNA end to G-rich motifs, which includes the 22 bp size of the motifs. Locations of APE1 within 92-156 bp range corresponds to the specific binding of the protein.

A similar analysis was done for complexes of APE1 with the control DNA substrate. The yield was comparatively low. The histrogam of the APE1 position measured from the end of the DNA molecule is shown in Figure S4A.

### 3.4. AFM data analysis: Sizes of DNA loops

For looped APE1-DNA complexes, two parameters were measured. In addition to the loop sizes, the lengths of the flanks were measured. The results are assembled in Figure 5. The loop sizes are shown in Fig. 5A and have a narrow distribution around 410 bp with a spread between ∼350 bp and ∼450 bp, which, when taking into account the 22 bp size of the G-rich motifs, corresponds to the assembly of complexes between the G-rich sites. Data beyond this size corresponds to the formation of non-specific loops.

**Figure 5.**
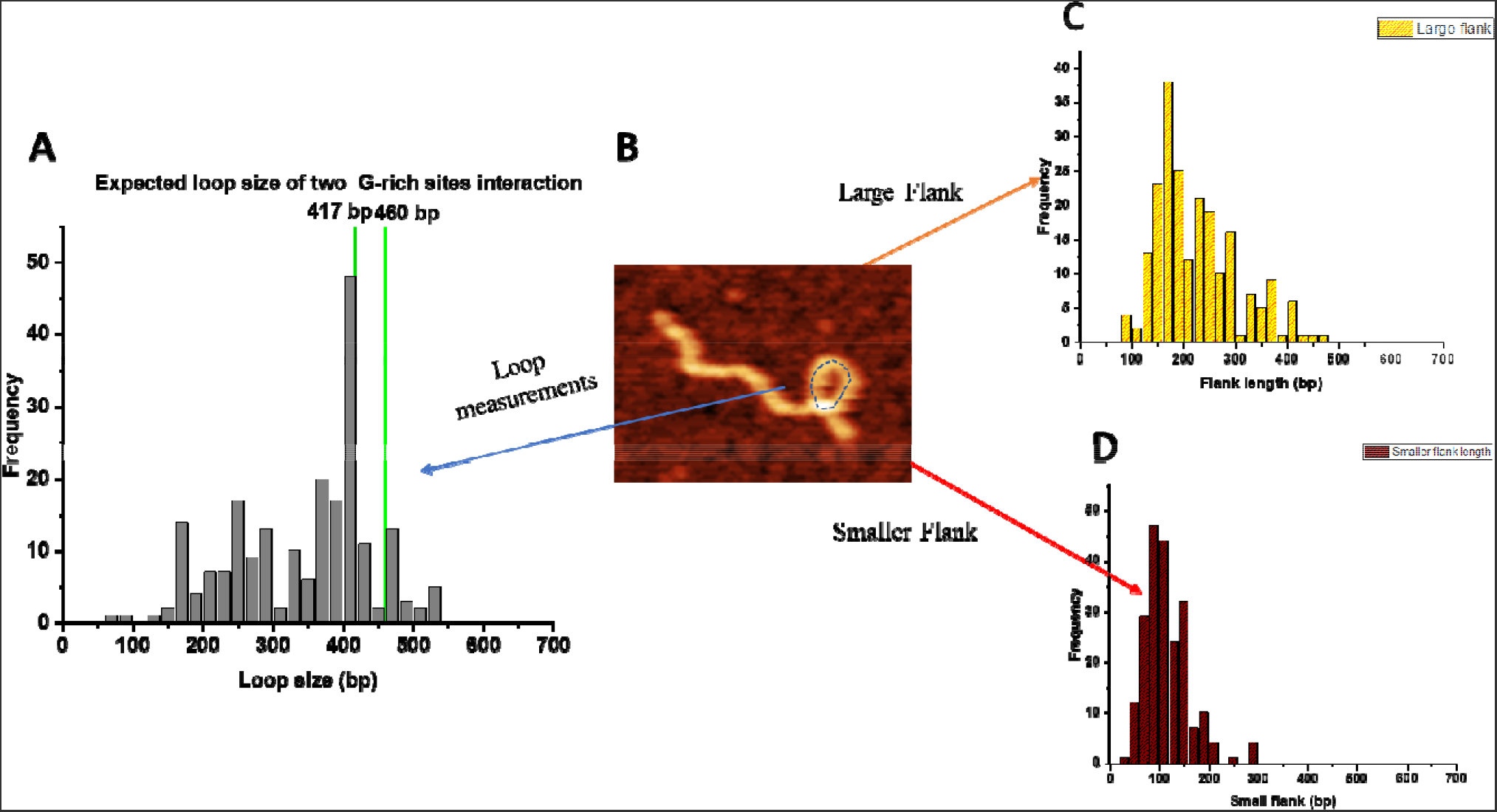
Looped complexes formed by APE1 on the G-rich DNA substrate. (A) The histogram for the loop sizes obtained for 200 looped complexes. Dotted vertical green lines indicate the sizes of loops formed by bridging of two G-rich motifs, which includes their sizes. (B) AFM image showing the looped complex. (C) The histogram of the lengths of the long arms. (D) The histogram of the lengths of short arms.

Results for measurements of short and long flanks are shown in Fig. 5C, D, respectively. The short flank length distribution is narrow and spans over the range of 60-150 bp, which due to the 22 bp length of the G-rich sites, covers the expected position for binding APE1 to one or the other G-rich site. On the other hand, the distribution of the lengths of the long arm is broad. In addition to the range corresponding to APE1 binding to one or the other G-rich site (vertical lines), events corresponding to the assembly of loops with APE1 binding to non-specific sites are also present.

The yield of looped complexes for the control DNA substrate was 4%, which is ∼1/6 compared to complexes assembled on G-rich DNA substrate (see Table 1). The control substrate’s loop sizes were analyzed, and the data are shown in Fig. S4B. The distribution was broad and flat, with no preferential loop size identifiable, indicative of a random distribution, which is corroborated by simulated distribution for non-soecific looping (Fig. S5).

### 3.5. Looped structures are formed by monomeric APE1

AFM captures the 3D shape of molecules and allows evaluation of their sizes. We used the height and volume measurements to estimate the molecular weights of proteins in complex with DNA (28,29). This information clarifies whether APE1 monomers, dimers, or larger oligomers are responsible for assembling loops. The two different pathways impose certain conditions; in the case of the dimeric stoichiometry in the looped complexes, each monomer should bind to DNA first, and then the loop is formed via protein-protein interactions. If the monomeric APE1 makes loops, the protein should have two DNA binding sites.

First, we measured the APE1 protein height in non-looped complexes with DNA on the G-rich substrate (Fig. 6A) and obtained the height histogram (Fig. 6B), which approximated with a Gaussian shows a peak at 1.2 ± 0.2 nm. The corresponding volume of the protein displayed a Gaussian distribution with a mean value of 125 ± 45 nm^3^ (Fig. 6C). Next, we measured the same parameters for APE1 in looped complexes (Fig. 6D). The height histogram for the protein in looped copmlexes was approximated with a Gaussian distribution and yielded a mean value of 1.1 ± 0.13 nm (Fig. 6E). Similarly, the volume of the protein displayed a Gaussian distribution centered around 130 ± 51 nm^3^ (Fig. 6F).

**Figure 6.**
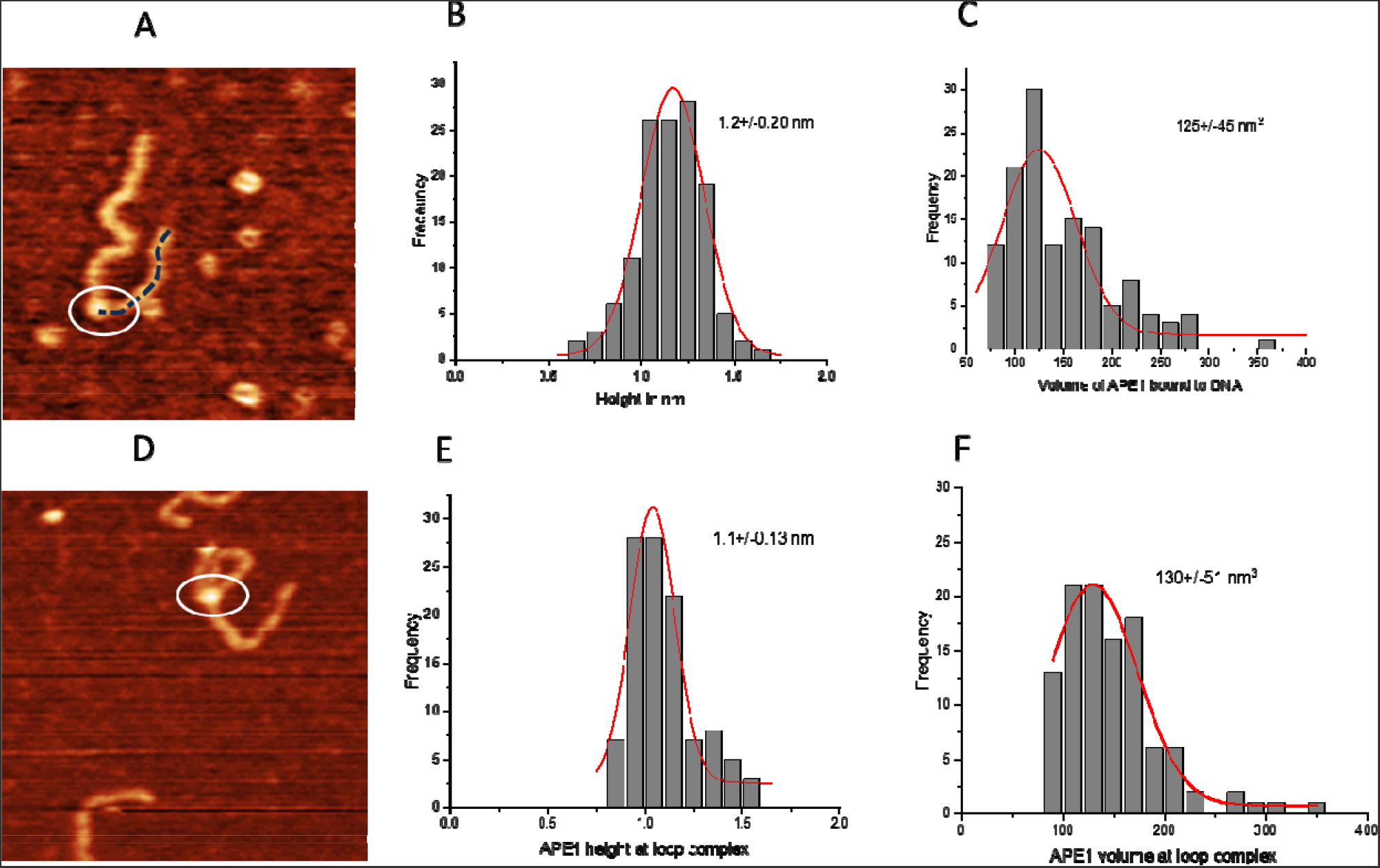
The height and volume analysis of the APE1 on G-rich DNA with the non-looped and looped complexes. (A) AFM image of the APE1 protein positioned on linear DNA. (B) Histograms for height values of the APE1 protein approximated with a Gaussian distribution (1.2 ± 0.20 nm). (C) The histogram of the volume measurements data approximated with a Gaussian distribution (125 ± 45 nm^3^). (D) AFM image of looped complexes of APE1 protein with DNA looped (circled in the. (F) The histogram for the protein height approximated with a Gaussian distribution (1.1 ± 0.13 nm). (D) The histogram for the protein volume approximated with a Gaussian distribution (130 ± 51 nm^3^).

As a control, we measured the height and volume of the APE1 protein in complexes with control DNA substrate. The height of the protein in this complex showed a Gaussian distribution with a mean value of the height 1.1 ± 0.13 nm, as shown in Fig. 7A. The volume of the protein exhibited a Gaussian distribution centered around 114 ± 19 nm^3^ (Fig. 7B). The height and volume of the APE1 protein in looped complexes with the control DNA produced values 1.07 ± 0.12 nm and 117 ± 14 nm^3^ (Fig. 7C, D), which are indistinguishable from those obtained for the G-rich DNA substrate.

**Figure 7.**
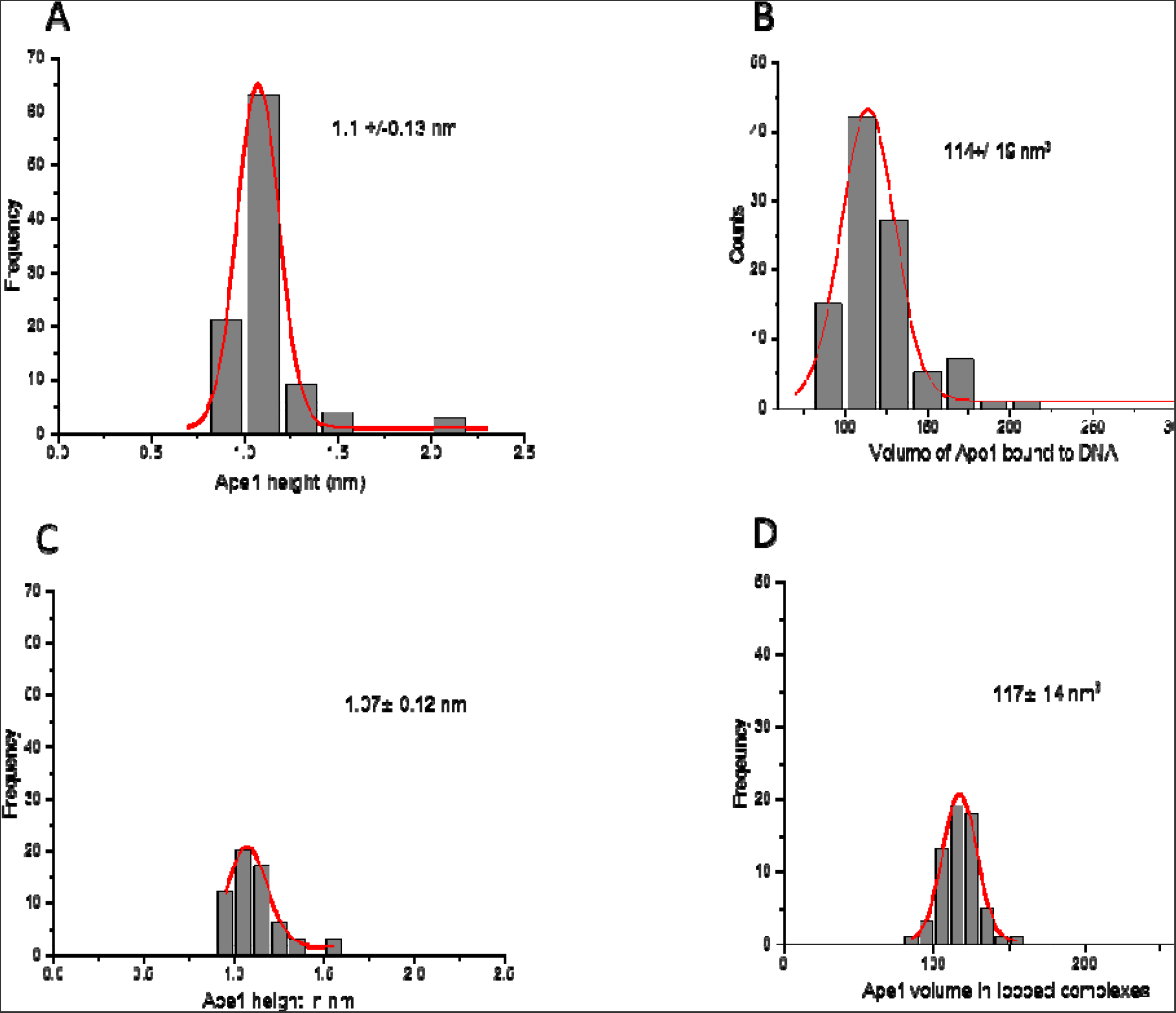
Height and volume measurements for complexes of the APE1 on control DNA substrate with unlooped and looped complexes. (A) and B are the histograms for the protein heights and volume, respectively for non-looped complexes. (C) and (D) are the histograms for the height and volume of APE1, respectively. Each histogram is approximated by single Gaussians with parameters indicated in the plots.

Height measurements of free APE1 produced the value 0.53 ± 0.14 nm (Fig. 8B), which, combined with the DNA height ∼ 0.5 nm, produces the height value ∼ 1.1. nm (Fig. 8C) for the protein bound to DNA. This value is close to the height values measured for protein bound to DNA, suggesting that protein binding to DNA is not accompanied by its oligomerization.

**Figure 8.**
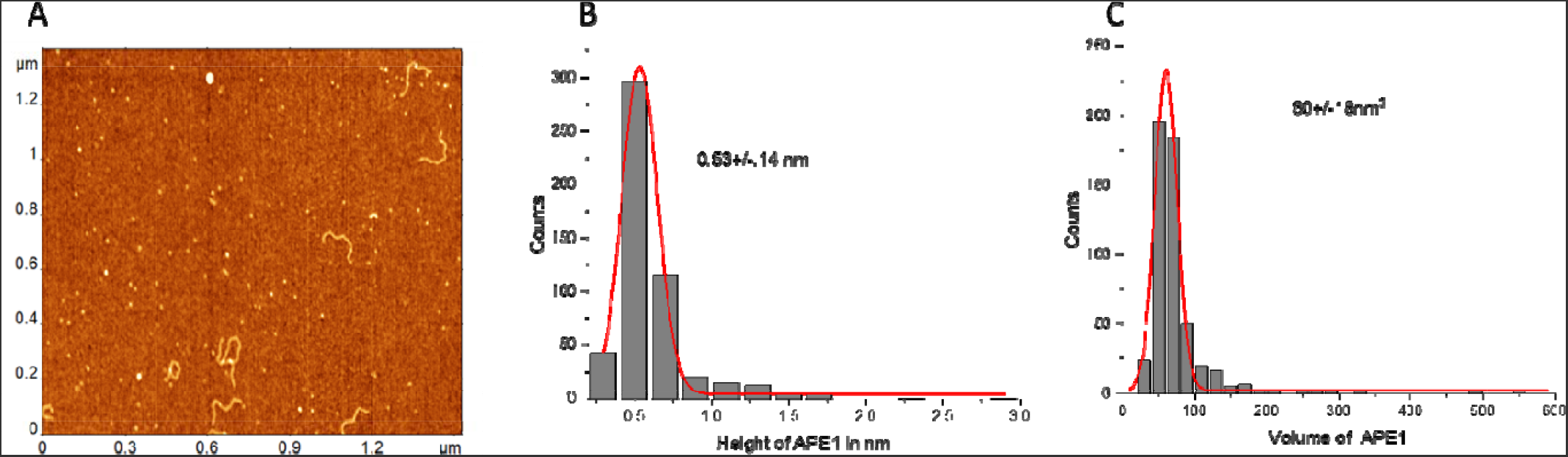
Height and volume measurements of the free APE1 protein. (A) AFM images of the free protein with added DNA as a reference. (B) and (C) are the histograms for the height and volume values built for 100 measurements. The histograms are approximated with Gaussians with parameters indicated in the plots.

These findings suggest that the monomeric form or single APE1 is involved in bridging two distant sites, suggesting that APE1 has two DNA binding segments and both are involved in the DNA looping.

## 4 Discussion

AFM studies clarified several novel features involved in the interaction of APE1 with DNA.

The affinity of APE1 to the G-rich motifs was hypothesized based on various indirect studies; here, direct visualization with AFM showed that such G-specificity exists. However, analysis of AFM data revealed that APE1 is capable of binding to non-G DNA as well, and the yield of such complexes is comparable with the formation of specific APE1-G complexes (Fig. 7).

Looping was another putative function of APE1 that we provide evidence for here. APE1 is capable of binding two sites on the same DNA molecule, leading to the formation of looped DNA structures. Analysis of AFM data showed that loops of different sizes are formed, and loops corresponding to the bridging of two G-rich segments were also visualized. The yield of such G-specific loops is close to the yield of non-specific loops, which is in line with the findings about the binding of APE1 to G-rich and non-specific linear DNA. However, additional quantitative analysis of looped complexes revealed an interesting assembly feature. As demonstrated in Figure 5, looped complexes in the vast majority of cases have short flanks with the length corresponding to the position of G-rich sites. In other words, APE1 in the loops bind to one G-rich site and one other site, which can be another G-rich site or any non-G rich segment.

The AFM images allowed us to elucidate the APE1 stoichiometry in the looped complexes. We determined the stoichiometry of APE1 in looped complexes by performing measurements of the protein sizes. The data shown in Figures 2, 3, and 4 demonstrate that looped complexes are formed by monomeric APE1 rather than its dimer. Bridging of two DNA binding sites is possible if the proteins has multiple DNA binding sites (25). Binding of two DNA segments by the monomeric APE1 suggests the protein has two binding sites. As we discussed above, looped complexes on the G-rich DNA substrate almost always have APE1 bound to one G-rich segment. This finding leads to the hypothesis that one DNA binding site of APE1 has a strong specificity to G-rich sequences, and the other site is more promicuous.

We have recently proposed the model for the site search process during DNA looping based on studies of the highly sequence-specific restriction enzyme SfiI (24,30). According to this model, during the search process, the protein initially binds to a specific site, grabs any non-specific site, and threads DNA in search of another specific site. In the framework of this model, we hypothesize that APE1 binds to the G-rich region on DNA at its specific site and searches DNA through using its less specific site. Note that such a mechanism has recently been proposed for the DNA looping for cohesin (31). APE1-mediated DNA looping for bringing two distant sites together may facilitate damage search in the transcriptional regulatory regions, coordinating repair and long-range promoter-enhancer interaction for repair and transcription.

Our data also shed light on whether APE1 recognizes G-rich regions (32). AFM images of the DNA templates (Fig. 1) demonstrate that the G-rich DNA segments are smooth and indistinguishable from the control. In contrast, G quadruplexes are considerably wider than the DNA duplex and can be visualized routinely with AFM (33,34). Thus, no quadruplexes are formed stably in the G-rich DNA substrate. At the same time, the specificity of APE1 to G-rich sequences was shown, suggesting that APE1 can bind to G-rich sequences without converting them into G quartet (14,35).

## Supplementary Materials

A detailed description of the DNA contour length, APE-1 positioning, and loop complexes measurements was provided.

## Author contributions

Y.L.L designed and supervised the project. K.K.B. initiated the project. S.V. and M.H. designed and performed the experiments and data analysis. Y.C. and S.P. prepared APE1 and DNA samples. All authors contributed to writing the manuscript.

## Note

The authors declare no competing financial interest.

## Acknowledgments

This work was supported by NSF grant MCB 1941049 to Y.L.L. We thank L. Shlyakhtenko (the University of Nebraska Medical Center) for valuable insights, Shaun Filliaux for proofreading the manuscript and all Y.L.L lab members for fruitful discussions of the results.

