## supplemental data for "Nanoscale interaction of endonuclease APE-1 with DNA characterized by atomic force microscopy"

*Corresponding authors

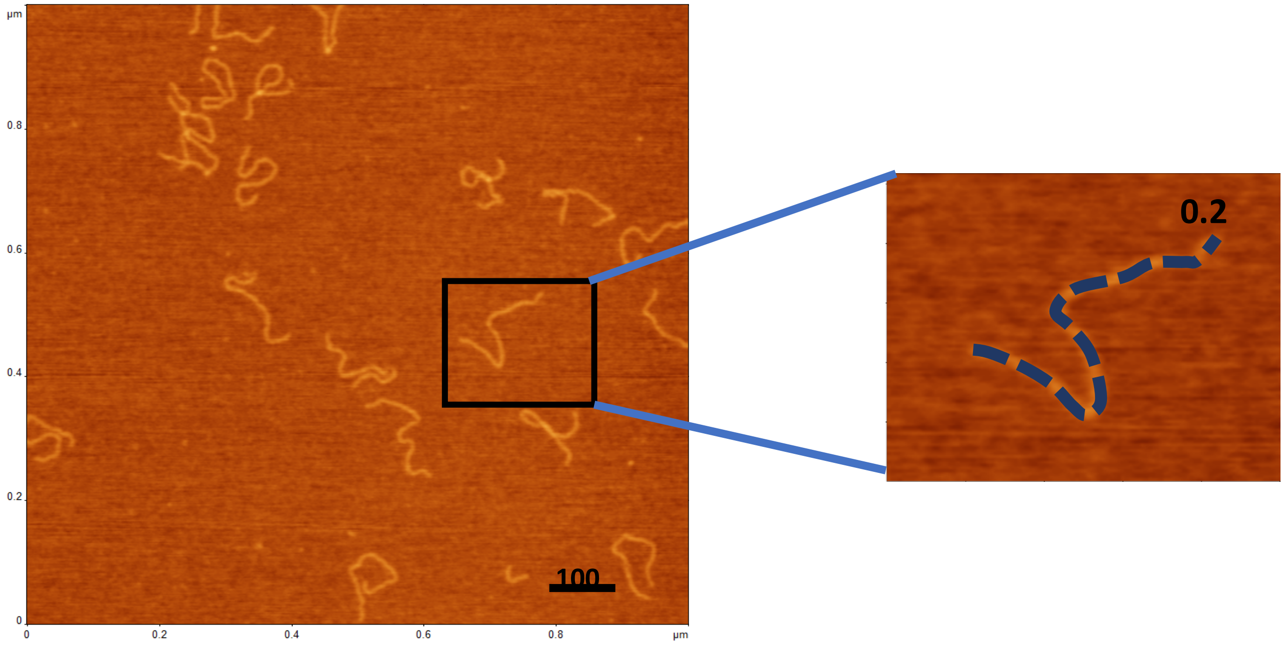


**Figure S1. DNA contour length measurement**. A typical 1x1 μm AFM scan for free DNA. All DNA filaments with non-crossed shape have a smooth morphology indicating no formation of G4 quadruplexes, which should appear as bright features on the AFM images (refs. 32, 33). The zoomed image of the DNA within the black box is shown to the right. The dotted line illustrates the procedure for the contour length measurements using FemtoScan software (Advanced Technologies Center, Moscow, Russia).

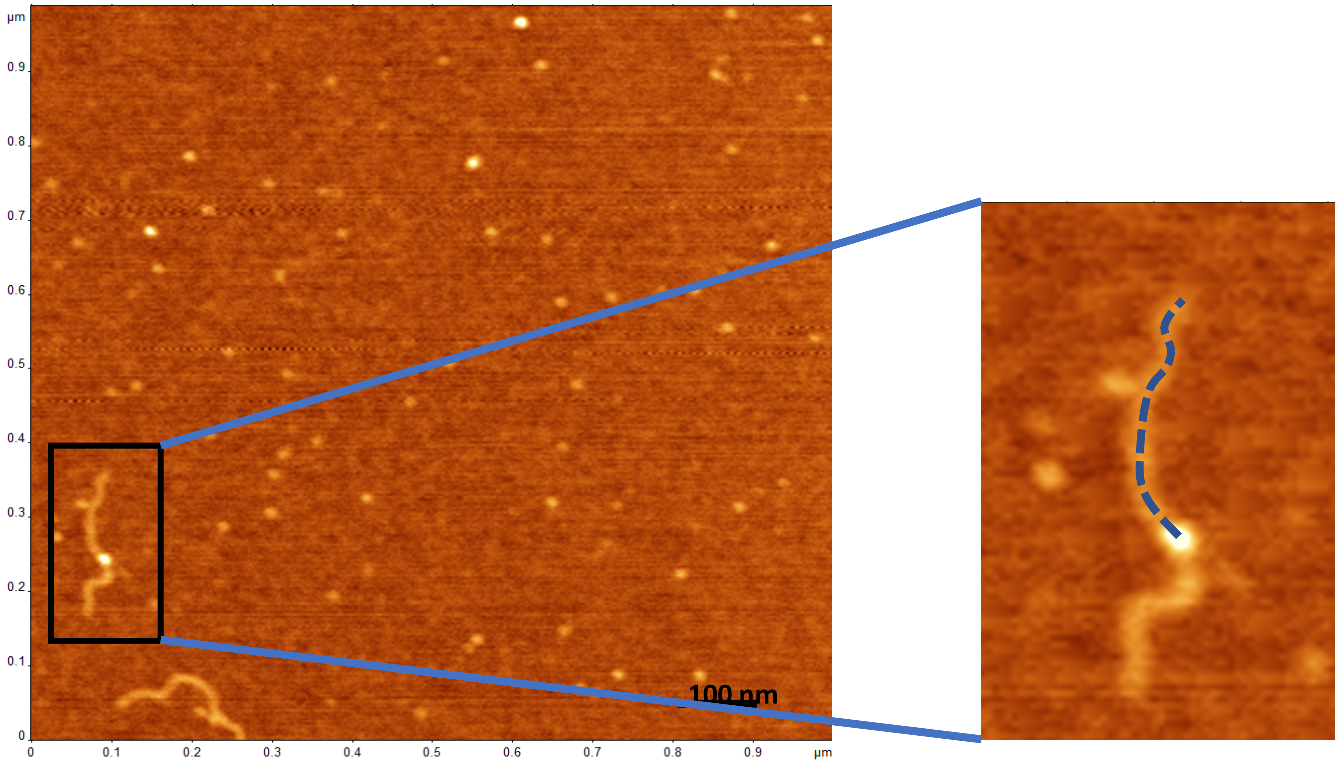


**Figure S2. AFM visualization of non-looped APE1-DNA complexes**. A typical 1x1 μm AFM image of complexes of APE1 with DNA. The zoomed image to the right illustrates the measurements of the length of one of the flanks. The measurements are made from the center of the bright feature to the end of the DNA. FemtoScan software (Advanced Technologies Center, Moscow, Russia) was used to perform the measurements.

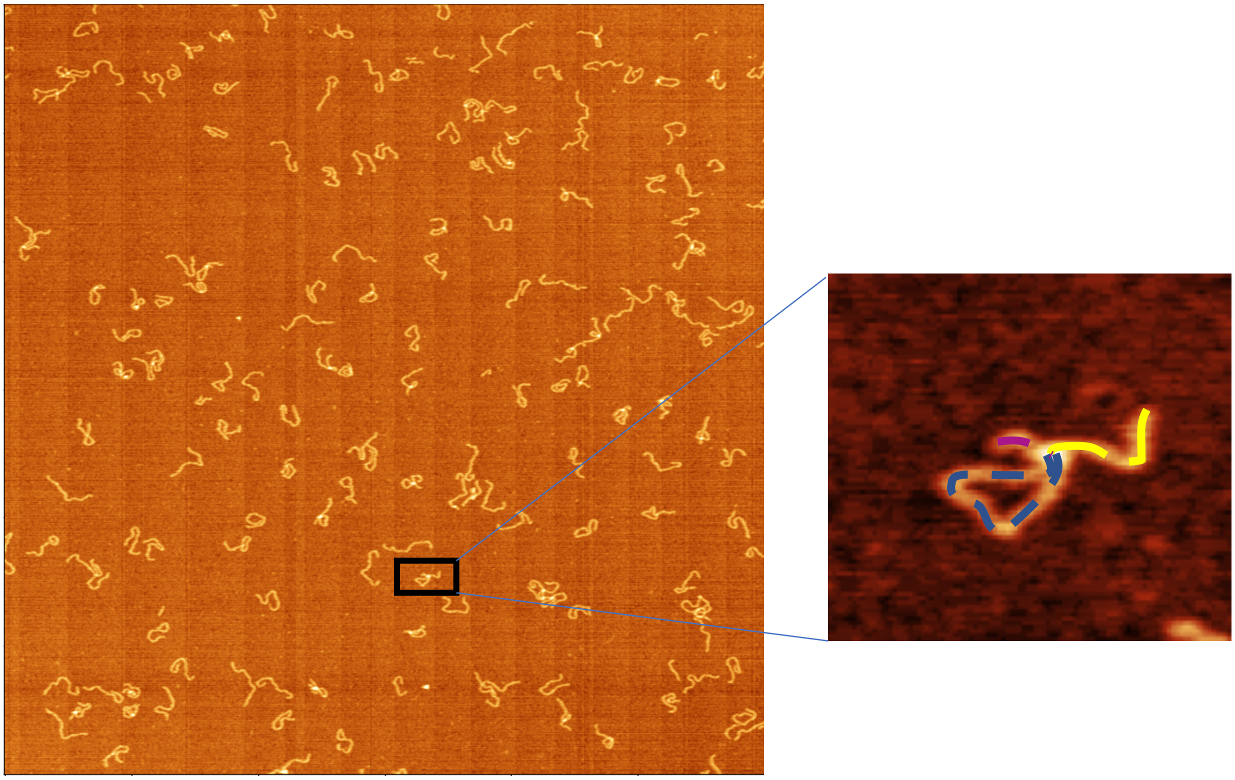

**Figure S3. APE1- DNA looped complexes**. Typical AFM images of APE1-DNA complexes for the AFM scan over 3x3 µm.The zoomed image to the right over the complex with the black box is shown to the right. The trace of the loop is indicated with a dotted line. The length of the DNA loop was measured starting from the center of the bright feature and ending at the same point. The lengths of the DNA flanks were measured from the end of DNA to the center of the bright feature. It is shown with the yellow line for the long flank and purple line for the short flank. Same procedure was followed to measure the loops of different sizes formed on the control DNA-APE-1 complexes. FemtoScan software (Advanced Technologies Center, Moscow, Russia) was used to perform the measurements.

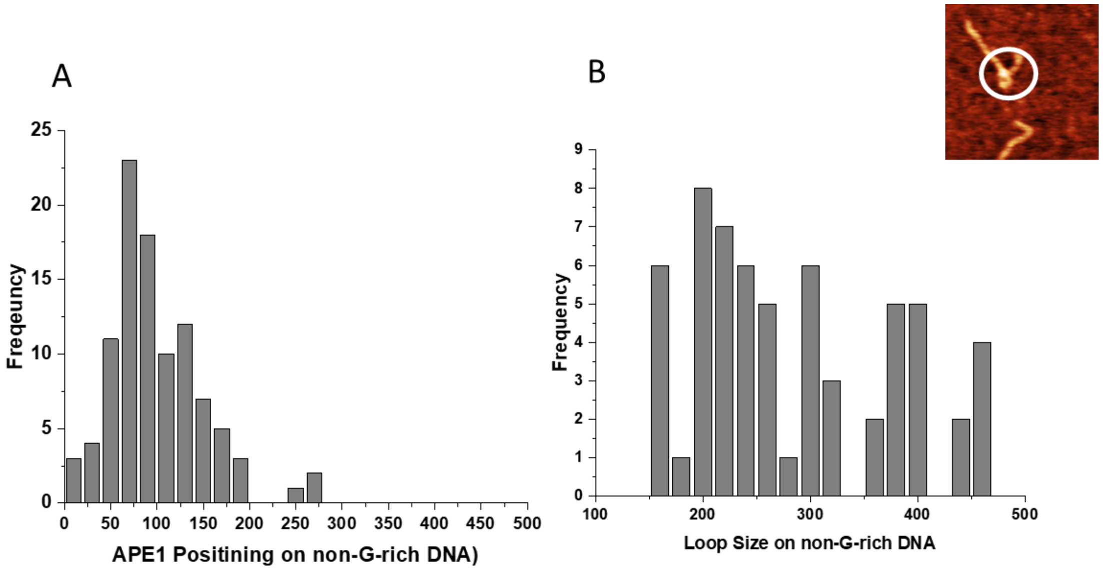


**Figure S4. Histograms of the APE1-DNA for the control DNA substrate.** (A) The histogram shows the positioning of APE1 on the non-G4 substrate. A total of N=100 complexes were measured and plotted. (B) The figure in the inset shows the loop size formed by the APE1-non-G4 DNA.A total of N=61 loop complexes were measured and plotted. With a bar graph showing a random loop size.

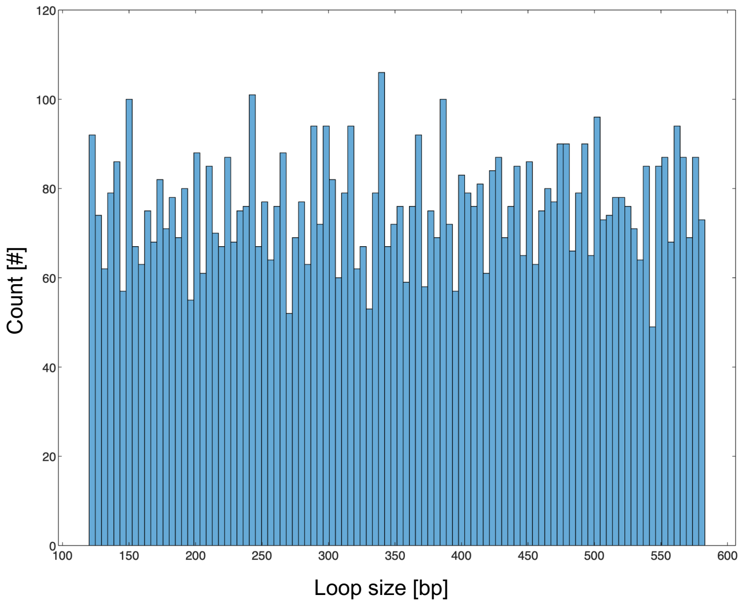


**Figure S5. Histogram of simulated random loop formation.** Monte Carlo simulations for random placement of APE1 was performed using bedtools to determine loop size distribution for non-specific loops.
